# Nup98 regulates the G1/S transition

**DOI:** 10.64898/2026.09.04.749412

**Authors:** Evi M. Malagise, Anna Sherman, Maya Capelson, Jared T. Nordman

## Abstract

Nuclear pore complexes (NPCs) are embedded throughout the nuclear envelope of all eukaryotic cells and are composed of 30 unique proteins called nucleoporins (Nups). While Nups primarily function at NPCs, multiple Nups function in the nucleoplasm independent of the NPC. We previously found that *Nup98-96* is necessary for entry into S phase in Drosophila cells, however, the mechanism is unknown. Here, we demonstrate that *Nup98-96* is one of only a subset of Nup genes that can promote entry into S phase. We identify that transcripts dependent on *Nup98-96* in Drosophila cells are enriched in cell cycle processes and that many of these transcripts are also dependent on the cell cycle regulator, E2f1. Importantly, we find that entry into S phase can be rescued by co-depleting the CycE/CDK2 inhibitor, *dacapo*, in *Nup98-96*-depleted cells. Furthermore, *Nup96-98* promotes S phase entry across multiple cell and tissue types and cell cycle contexts. Critically, overexpression of *Nup98* has a dominant-negative effect on entry into S phase in larval salivary glands, suggesting that Nup98, not Nup96, promotes entry into S phase. Our work identifies Nup98 as a key regulator of the G1/S transition in metazoans.

## INTRODUCTION

All eukaryotic cells store genetic material inside the boundary of the double-membrane nuclear envelope, which is permeated by large protein assemblies called nuclear pore complexes (NPCs) that facilitate nucleocytoplasmic transport. NPCs are composed of multiple copies of 30 unique proteins called nucleoporins (Nups). There are two major classifications of Nups: dynamic Nups and core scaffold Nups. Dynamic Nups can shuttle on and off pores and function within the nucleoplasm, whereas core scaffold Nups are stably associated with NPCs [1–3]. A subset of Nups can bind to chromatin, alter chromatin condensation, and affect transcriptional activity [4–11]. Two striking examples of Nup functional diversity are Nup98 and Nup96. Nup98 and Nup96 are produced from the *Nup98-96* gene locus, which is transcribed, translated, and folded as a single polyprotein product that is then cleaved into Nup98 and Nup96 via autoproteolysis [12–14]. While Nup96 is a core scaffold Nup that functions as part of the Nup107-160 subcomplex, Nup98 is a dynamic phenylalanine-glycine (FG) Nup with a unique glycine-leucine-phenylalanine-glycine (GLFG) repeat domain that binds to chromatin at active genes, regulates transcriptional activity of developmental and cell cycle genes, influences enhancer-promoter contacts, regulates transcriptional memory, and functions during heterochromatin repair responses in Drosophila [1–6,8,9,15–18]. Nup98 also localizes to enhancers and promoters and affects transcription in mammalian cells [19–21]. These intranuclear and transcriptional functions of Nup98 are particularly interesting, as chromosomal rearrangements involving *Nup98* result in gene fusions in humans. These fusions generate chimeric oncoproteins that are implicated in the pathogenesis of multiple hematopoietic malignancies [22,23]. The GLFG domain of Nup98 is critical for Nup98-oncofusion protein functions in hematopoietic malignancies [23–28].

Previously, we demonstrated that *Nup98-96* is necessary to promote S phase entry in Drosophila [29]. The mechanism by which *Nup98-96* regulates entry into S phase, however, was not explored. Progression through the G1/S transition is tightly regulated by the E2F family of transcription factors. While mammals contain multiple E2F activators and repressors [30,31], *D. melanogaster* contains a single activator and repressor (E2f1 and E2f2, respectively) [30–32]. E2f1 activates transcription of cell cycle and S phase genes and a critical E2f1 target gene is *CycE*, which is crucial for the G1/S transition [33–37]. In Drosophila, CycE/CDK2 are the sole cyclin/CDK complex needed for the G1/S transition, as CycD/CDK4 are required for growth but not the G1/S transition [38,39]. Despite the known regulators of the G1/S transition, it is likely that there are other factors that function during entry into S phase.

In this study, we investigate how *Nup98-96* promotes S phase entry. We find that *Nup98-96* is one of only a subset of Nup genes that regulates S phase entry, and this activity is likely independent of NPC function. *Nup98-96* regulates the abundance of key cell cycle and S phase gene transcripts, many of which are canonical E2f1 targets. Depleting *Nup98-96* or *E2f1* causes reduced S phase entry that can be suppressed by co-depleting the CycE/CDK inhibitor, Dacapo [40,41]. *Nup98-96* is required for S phase entry in multiple developmental and cell cycle contexts and overexpression of *Nup98* phenocopies *Nup98-96* depletion in the larval salivary gland. Taken together, our findings provide insight into how the G1/S transition is regulated and define a function for Nup98 at the G1/S transition.

## RESULTS

### A subset of Nups impact the G1/S transition

In Drosophila, depletion of *Nup98-96* causes a reduction in cells in S phase and an increase in cells in G1 phase [29]. This phenotype suggests that *Nup98-96* functions in the entry into S phase. It is not known, however, if this phenotype is specific to *Nup98-96* or whether this is a common phenotype upon depletion of individual Nups or the NPC as a whole. To test this directly, we depleted each of the 30 metazoan Nups by RNA interference (RNAi) in Drosophila S2 cultured cells and measured the fraction of cells in each cell cycle phase based on DNA content and 5-Ethynyl-2’-deoxyuridine (EdU) incorporation after three or five days of RNAi depletion (**Figure 1A-C**; **Supplemental Figure 1A-C**). All depletions were verified by qPCR or western blotting (**Supplemental Figure 1D-F**). If disruption of NPC function non-specifically affects the G1/S transition, we would predict the majority of Nup depletions would reduce the fraction of cells in S phase with a concomitant increase in cells in G1 phase. If, on the other hand, individual Nups affect the G1/S transition independently of the NPC, we expect only a subset of Nups to promote S phase entry.

**Figure 1:**
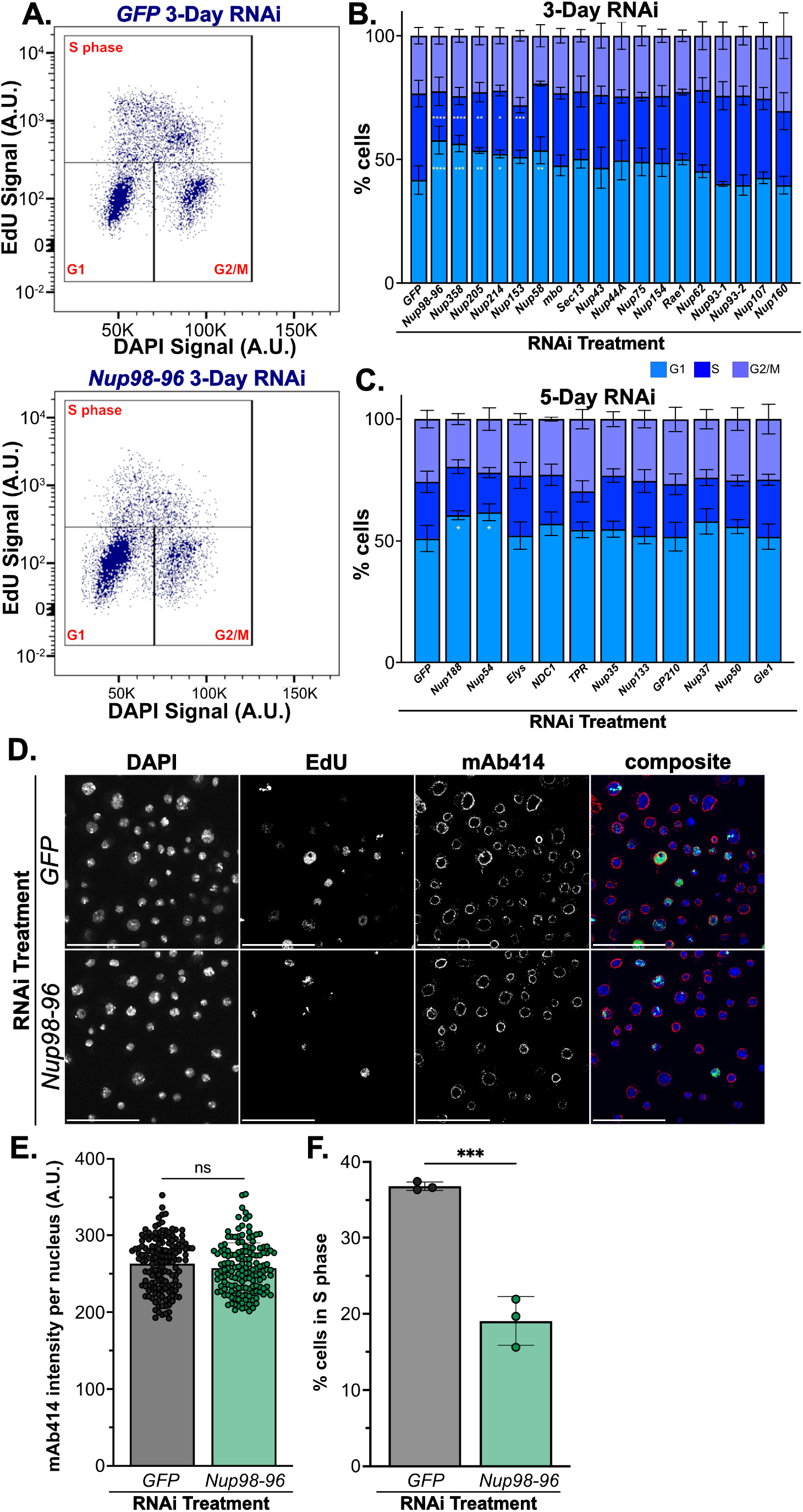
A subset of Nups impacts the G1/S transition. **(A)** Horseshoe plots were generated by plotting DNA content (DAPI) against EdU signal. Representative plots from 3-day control RNAi (*GFP* RNAi) and 3-day *Nup98-96* RNAi cells are shown. Gates indicating G1, S, and G2/M populations are shown (black boxes) with cell cycle phase noted (red). **(B-C)** RNAi was conducted for either 3 days **(B)** or 5 days **(C)** targeting control and Nups. The percentages of cells in G1 (light blue), S (blue), and G2/M (purple) phases were quantified. Statistical significance was determined via One-Way ANOVA with *post-hoc* Dunnett’s test (*GFP* RNAi as control; n=3-9; * p<0.05, ** p<0.01, *** p<0.001, **** p<0.0001). Error bars represent SD. **(D)** Representative images of non-targeting *GFP* and *Nup98-96* RNAi cells that were pulsed with EdU (green), stained with mAb414 (red), and stained with DAPI (blue). Scale bars = 50 μm. **(E)** mean mAb414 intensity per nucleus was quantified for 50 nuclei per replicate (n=3). Statistical significance was determined via Student’s T-test. Error bars represent SD. **(F)** The percentage of cells in S phase was quantified by measuring the EdU intensity per nucleus for 753-1177 nuclei per replicate and separating cells into EdU-positive and EdU-negative subgroups (n=3). Error bars represent SD. Statistical significance was determined via Student’s T-test. *** p<0.001.

After individually depleting all 30 Nups by RNAi, we found that depletion of only four (*Nup98-96*, *Nup358*, *Nup205*, and *Nup214)* affected the G1/S transition. This suggests that a general disruption of NPC function does not impact entry into S phase (**Figure 1B-C**). These Nups have different localizations and functions within NPCs. Nup358 and Nup214 are cytoplasmic filament FG Nups, Nup98 is a central FG Nup, Nup96 is an outer ring Nup, and Nup205 is an inner ring Nup [2,42,43]. Thus, it is likely that depletion of *Nup98-96*, *Nup358*, *Nup205* or *Nup214* impact S phase entry through different mechanisms. Interestingly, of the Nups that display a reduced entry into S phase phenotype, only Nup98 is known to be a dynamically associated with the NPC, meaning it can shuttle on- and off-pore and function within the nucleoplasm [1–3,43]. Therefore, we chose to focus on how *Nup98-96* functions to regulate the G1/S transition.

To ensure that the *Nup98-96* depletion phenotype was specific to *Nup98-96* depletion and not an off-target effect, we validated our results with two additional independent interfering RNAs (**Supplemental Figure G-H**). To test the possibility that depletion of *Nup98-96* indirectly affects the G1/S transition by reducing the abundance of nuclear pores, we measured the relative abundance of NPCs in control and *Nup98-96*-depleted cells. We performed confocal microscopy using the mAb414 antibody that binds to FG repeats within Nups as a marker of NPC abundance [44]. These same samples were pulsed with EdU to count the fraction of cells in S phase. Similar to previous studies in Drosophila larval salivary glands and imaginal discs [18], we did not detect a significant decrease in the mean mAb414 intensity between *Nup98-96* and control depletions (**Figure 1D-E**). Critically, we confirmed that depletion of *Nup98-96* reduced the percentage of cells in S phase relative to control cells in mAb414-stained cells with EdU labeling (**Figure 1F**). Taken together, our data suggests that depletion of *Nup98-96* prevents cells from entering S phase without destabilizing NPCs.

### *Nup98-96* regulates the transcriptional activation of cell cycle-related genes and E2f target genes

Given the transcriptional functions of Nup98, we asked if reduced S phase entry upon *Nup98-96* depletion could be explained by changes in gene expression. To test this, we measured changes in global mRNA levels after three days of *Nup98-96* depletion relative to control depletions by RNA-sequencing (RNA-seq). We confirmed that samples cluster by RNAi target (**Supplemental Figure 2A-B**). After depletion of *Nup98-96*, there were 551 significantly upregulated and 440 significantly downregulated transcripts (padj <0.05) (**Figure 2A**). We conducted Gene Ontology (GO) analyses on significantly upregulated and downregulated transcripts (padj <0.05) to identify processes perturbed upon *Nup98-96* depletion (**Figure 2B-C**). Transcripts that were upregulated after *Nup98-96* depletion were most significantly enriched in general processes, including *cellular process* (FDR 3.11×10^-12^), *response to stimulus* (FDR 2.28×10^-8^), and *animal organ morphogenesis* (FDR 8.16×10^-6^) (**Figure 2B**). Given that these transcripts are not associated with cell cycle regulation, this suggests that the reduced entry into S phase observed upon *Nup98-96* depletion is not caused by increased transcript levels. We analyzed the significantly downregulated transcripts (padj <0.05) and found that six of the top ten most enriched biological processes were related to the cell cycle (**Figure 2C**). This includes the top three categories of *cell cycle* (FDR 8.0×10^-15^), *mitotic cell cycle* (FDR 1.08×10^-13^) and *cell cycle process* (FDR 1.35×10^-12^) (**Figure 2C**). Many of the downregulated genes are critical for cell cycle progression, including *cyclins*, *CDKs*, and *Dp* [31]. Interestingly, many of the genes downregulated after *Nup98-96* depletion are canonical targets of the E2f1 transcription factor that promotes S phase entry [34,35].

**Figure 2:**
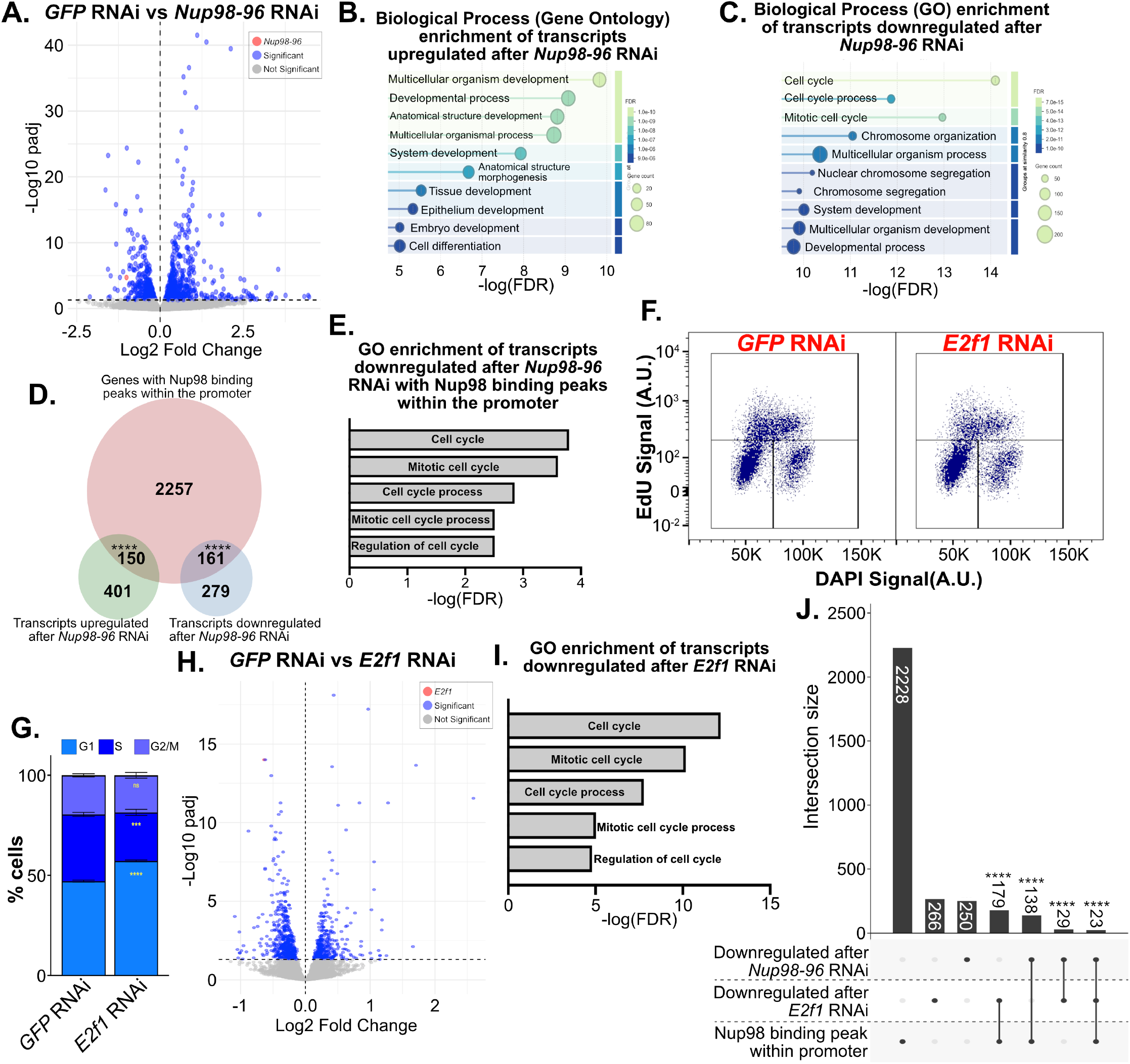
Nup98-96 regulates the transcriptional activation of cell cycle-related genes and E2f target genes. **(A)** Volcano plot was generated by plotting -Log10(padj) against Log2 fold change. Points represent transcripts with no significant changes (gray), transcripts with significant changes (blue), and *Nup98-96* (red). **(B)** Gene ontology analysis of significantly upregulated transcripts after *Nup98-96* depletion sorted by false discovery rate (FDR). Color scale indicates FDR and circle size represents transcript count. **(C)** Gene ontology analysis of significantly downregulated transcripts after *Nup98-96* depletion sorted by FDR. Color scale indicates FDR and circle size represents transcript count. **(D)** Venn diagram represents the overlap between genes that had Nup98 binding peaks with the promoter region (red), are upregulated after *Nup98-96* RNAi (green), and downregulated after *Nup98-96* RNAi (blue). Numbers represent transcript count. Statistical significance was determined via hypergeometric test. **** p<0.0001. **(E)** Gene ontology analysis of transcripts that are significantly downregulated after *Nup98-96* depletion and contain Nup98 binding peaks within the promoter region. x-axis represents –log(FDR). **(F)** Horseshoe plots were generated by plotting DNA content (DAPI) against EdU signal. Representative plots from 3-day control RNAi (*GFP* RNAi) and 3-day *E2f1* RNAi cells are shown. RNAi targets are noted in red. Gates indicating G1, S, and G2/M populations are shown (black boxes). **(G)** The percentages of cells in G1 (light blue), S (blue), and G2/M (purple) phases were quantified and statistical significance was determined via Student’s T-test (GFP as control; n=3; **** p<0.0001; *** p<0.001; ns=not significant). Error bars represent SD. **(H)** Volcano plot was generated by plotting -Log10(padj) against Log2 fold change. Points represent transcripts with no significant changes (gray), transcripts with significant changes (blue), and *E2f1* (red). **(I)** Gene ontology analysis of significantly downregulated transcripts after *E2f1* depletion. x-axis represents –log(FDR). **(J)** Upset plot represents overlap between transcripts downregulated after *Nup98-96* RNAi, transcripts downregulated after *E2f1* RNAi, and genes with Nup98 binding peaks at the promoter. Numbers in/above each bar indicate the number of unique genes within that set. Statistical significance was determined via multivariate hypergeometric test. **** p<0.0001.

Nup98 functions independently of the nuclear pore to regulate transcription [5]. To probe whether the differentially expressed transcripts could be regulated by Nup98 directly, we asked what fraction of differentially expressed transcripts after *Nup98-96* depletion contain Nup98 binding sites in their promoter regions based on published Nup98 chromatin immunoprecipitation sequencing (ChIP-seq) data [9]. We found that 27.2% of upregulated transcripts and 36.6% of downregulated transcripts also had Nup98 binding peaks specifically within the promoter region, which is much higher than expected by chance (p<0.0001, hypergeometric test; **Figure 2D**). Genes that contain Nup98 binding peaks within their promoter region and produce transcripts dependent on *Nup98-96* were significantly enriched in the cell cycle-related biological processes, including *cell cycle* (FDR 1.6×10^-4^), *mitotic cell cycle* (FDR 2.5×10^-4^), *cell cycle process* (FDR 1.4×10^-3^), *mitotic cell cycle process* (FDR 3.1×10^-3^), and *regulation of cell cycle* (FDR 3.1×10^-3^) (**Figure 2E**). Taken together, these data suggest that the reduced entry into S phase observed upon *Nup98-96* depletion may be caused by deficits in Nup98-dependent transcriptional activity.

In Drosophila S2 and Kc cells, Nup98 ChIP-seq peaks are enriched in DNA Replication-related Element Factor (DREF) binding motifs (**Supplemental Figure 2C-D**) [8,9,45]. DREF is a transcriptional regulator of DNA replication- and growth-related genes [46–48] and of *E2f1* [49,50]. DREF motifs are enriched in binding sites of the DREF transcription factor [46]. Thus, Nup98 may work together with either DREF or E2f1 to control expression of genes involved in cell cycle regulation. To address this possibility, we compared the transcriptional profiles of S2 cells depleted for *DREF* or *E2f1* with *Nup98-96*-depleted cells. We analyzed published RNA-seq data from *DREF*-depleted and control non-targeting *lacZ*-depleted S2 cells (GEO: GSE117217; [51]) and found that transcripts downregulated after *DREF* depletion are not enriched in similar GO processes as *Nup98-96-*depleted cells (**Supplemental Figure 2E**).

In contrast, the changes in transcript levels and reduced percentage of cells entering S phase upon *Nup98-96* depletion are similar to reported phenotypes associated with reduced E2f1 function [35]. To directly compare these transcriptional similarities, we depleted *E2f1* for three days and measured the percentage of cells in each cell cycle phase and global changes in mRNA levels. Similar to depleting *Nup98-96*, depletion of *E2f1* for three days caused a decrease in S phase cells with a concomitant increase in G1 cells (G1 p<0.0001; S p<0.001; **Figure 2G**). Next, we measured changes in global mRNA transcript levels by RNA-seq analysis and confirmed that samples cluster by RNAi target (**Supplemental Figure 2G-F**). After depletion of *E2f1*, there were 346 significantly upregulated and 497 significantly downregulated transcripts (padj <0.05; **Figure 2H**). GO analysis on the significantly upregulated transcripts (padj <0.05) revealed non-specific GO categories with *cellular process* being the most significant (FDR 2.25×10^-9^; **Supplemental Figure 2H**). GO analysis of downregulated transcripts (padj <0.05) revealed terms associated with translation and biogenesis (**Supplemental Figure 2I**). *Cell cycle* (6.81×10^-13^), *mitotic cell cycle* (6.65×10^-11^), *cell cycle process* (1.72×10^-8^), and other cell cycle-related processes were also enriched in the transcripts that are dependent on E2f1 for expression (**Figure 2I**). When comparing the 440 transcripts that were downregulated upon *Nup98-96* depletion and the 497 transcripts that were downregulated upon *E2f1* depletion, 52 are shared, which is more than expected by chance (p <0.0001, hypergeometric test; **Supplemental Figure 2J, Supplemental Table 1**). We then asked if there are transcripts that are downregulated upon *Nup98-96* depletion and *E2f1* depletion that also contain Nup98 binding peaks within their promoter region. We identified 23 genes that meet these criteria, which is more than expected by chance (p<0.0001, multivariate hypergeometric test; **Figure 2J; Supplemental Table 2**). Critically, the primary cyclin responsible for propelling cells through the G1/S transition, *cycE*, was downregulated after both *Nup98-96* depletion and *E2f1* depletion and contains Nup98 binding peaks within its promoter region (**Supplemental Table 2**). Taken together, our data indicates that *Nup98-96* affects the regulation of many important cell cycle genes and shares cell cycle and transcriptional phenotypes that are similar to cells depleted of *E2f1*.

### *Nup98-96* and *dacapo* have opposing functions in regulating the G1/S phase transition

Cells depleted of *Nup98-96* have cell cycle progression and transcriptional changes that are similar to cells depleted of *E2f1* (**Figure 2**). This raises the possibility that Nup98 or Nup96 controls S phase entry. If so, we would expect that co-depletion of a negative regulator of S phase entry could suppress the reduced S phase entry upon Nup98-96 depletion. To test this possibility, we co-depleted *Retinoblastoma factor* (*Rbf*) or *dacapo* (*dap*) in *Nup98-96*-depleted cells. Rbf inhibits E2f1 and E2f-dependent transcription [52] while Dap directly inhibits CycE/CDK2 [40,41]. Depletions were confirmed by qPCR or western blotting (**Supplemental Figure 3A-C**). We first singly depleted *Rbf*, *dap*, *Nup98-96* or non-targeting *GFP* and measured the percentage of cells in each cell cycle phase by flow cytometry (**Figure 3A-B**). We saw no significant changes in the percentage of cells in any phase when *Rbf* or *dap* were depleted alone relative to control cells (**Figure 3B**). Consistent with our previous results, we observed an increase in G1 cells and a decrease in S phase cells when *Nup98-96* was depleted (G1 p<0.001; S p<0.01; **Figure 3B**). Co-depletion of *Rbf* in *Nup98-96*-depleted cells did not rescue entry into S phase, as there was still an increase in G1 cells and a decrease in S phase cells (G1 p<0.001; S p<0.05; **Figure 3B**). Interestingly, co-depletion of *dap* in *Nup98-96*-depleted cells fully rescued the entry into S phase (**Figure 3B**). To validate these findings, we repeated the co-depletions of *dap* and *Nup98-96* and measured the percentage of S phase cells by fluorescent microscopy in EdU-pulsed cells. As expected, we saw a significant decrease in S phase cells when *Nup98-96* was depleted (p<0.05) and a rescue in the percentage of S phase cells when *dap* and *Nup98-96* were co-depleted (**Figure 3C-D**). Taken together, our data indicate that *dap* and *Nup98-96* have opposing functions in regulating the G1/S phase transition.

**Figure 3:**
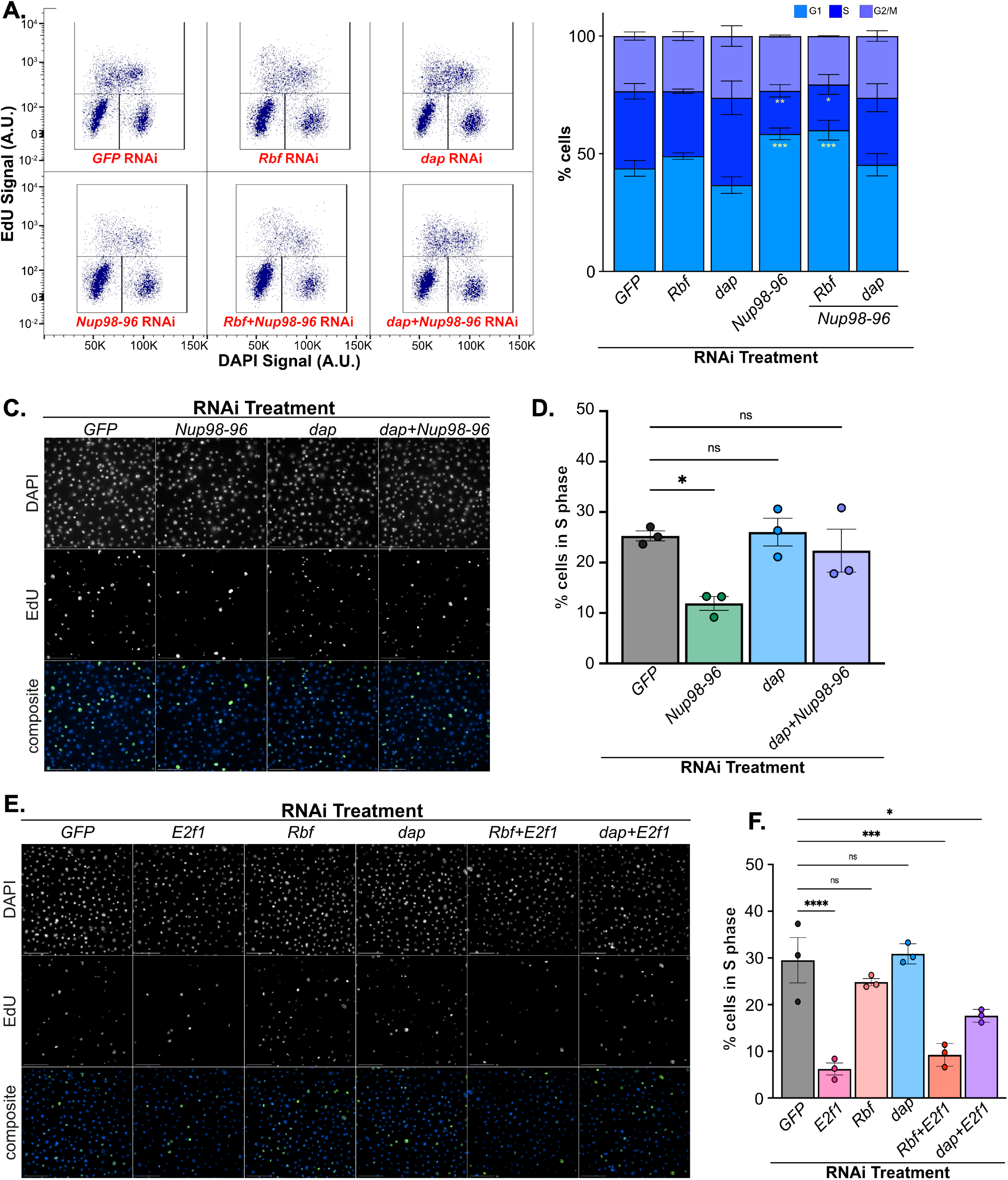
Nup98-96 and Dacapo have opposing functions in regulating the G1/S transition. **(A)** Horseshoe plots were generated by plotting DNA content (DAPI) against EdU signal. Representative plots from control *GFP* RNAi, 3-day *Nup98-96* RNAi, 5-day *Rbf* RNAi, 5-day *dap* RNAi, *Rbf+Nup98-96* dual RNAi, and *dap+Nup98-96* dual RNAi cells are shown. Gates indicating G1, S, and G2/M populations are shown (black boxes). RNAi targets are noted in red. **(B)** The percentages of cells in G1 (light blue), S (blue), and G2/M (purple) phases were quantified and statistical significance was determined via One-Way ANOVA with *post-hoc* Dunnett’s test (*GFP* RNAi as control; n=3; *** p<0.001, ** p<0.01, * p<0.05). Error bars represent SD. **(C)** Representative images of control (*GFP*), *Nup98-96*, *dap*, and dual *dap+Nup98-96 RNAi* cells that were pulsed with EdU (green) and stained with DAPI (blue). Scale bars = 50 μm. **(D)** The percentage of cells in S phase was quantified by measuring the mean EdU intensity per nucleus for 502-1929 nuclei per replicate and separating cells into EdU-positive and EdU-negative subgroups. Statistical significance was determined via One-Way ANOVA with *post-hoc* Dunnett’s test (*GFP* RNAi as control; n=3; * p<0.05, ns=not significant). **(E)** Representative images of control (*GFP*), *E2f1*, *Rbf*, *dap*, *Rbf+E2f1*, and *dap+E2f1* RNAi cells were pulsed with EdU (green) and stained with DAPI (blue). Scale bars = 50 μm. **(F)** The percentage of cells in S phase was quantified by measuring the mean EdU intensity per nucleus for 901-1536 nuclei per replicate and separating cells into EdU-positive and EdU-negative subgroups. Statistical significance was determined via One-Way ANOVA with *post-hoc* Dunnett’s test (*GFP* RNAi as control; n=3; **** p<0.0001; *** p<0.001; * p<0.05, ns=not significant).

To further test the possibility that *Nup98-96* functions similarly to *E2f1* during entry into S phase, we singly depleted *E2f1*, *dap*, *Rbf*, or non-targeting *GFP* and measured the percentage of cells in S phase by fluorescent microscopy (**Figure 3E-F**). As expected, *E2f1*-depleted cells had a significant reduction in the percentage of cells in S phase (p<0.0001; **Figure 3F**). Similar to depleting *Rbf* in *Nup98-96*-depleted cells, depletion of *Rbf* in *E2f1*-depleted cells did not rescue entry into S phase (p<0.001; **Figure 3F**). Entry into S phase was partially rescued, however, when *dap* was depleted in *E2f1*-depleted cells (p<0.01; **Figure 3F**). Considering the necessity of E2f1 transactivation of *CycE* for entering S-phase [33,34,36], it is possible that increased CycE/CDK2 activity after *dap* co-depletion allowed more of the cell population to enter S phase but not to the level of control. Given that depletions of *Nup98-96* and *E2f1* result in similar phenotypes, that *E2f1* and *dap* have known opposing functions during S phase entry, and that co-depletion of *dap* and *Nup98-96* rescues entry into S phase, we conclude that *Nup98-96* and *dap* have opposing functions during the G1/S transition.

### Nup98 regulates entry into S phase in Drosophila salivary glands

While *Nup98-96* controls entry into S phase in Drosophila cultured cells, we wanted to test if *Nup98-96* is required for S phase entry in Drosophila tissues. We chose to test this in Drosophila larval salivary glands, as salivary gland cells undergo endoreplication, repeated S-G cycles in the absence of mitotic divisions [53,54]. In larval salivary glands, tissue growth is directly tied to ploidy [55,56]. Thus, the size of salivary glands provides a direct readout of cell cycle progression. To test whether *Nup98-96* regulates entry into S phase in larval salivary glands, we depleted *Nup98-96* in 96-99hr after egg laying (AEL) salivary glands using the *fkh*>GAL4 driver and measured both ploidy and nuclear volume. First, we observed a decrease in tissue size in *Nup98-96-*depleted salivary glands relative to *fkh*>GAL4 control salivary glands (**Supplemental Figure 4A**). This size difference was not due to fewer cells, as the number of nuclei in *Nup98-96 RNAi* and control salivary glands were not significantly different (**Supplemental Figure 4B**).

To assess relative ploidy of salivary gland nuclei, we measured the total DAPI content of individual nuclei for *Nup98-96*-depleted, *dap*-depleted, and *dap- and Nup98-96*-depleted salivary glands relative to control glands that contain the *fkh*>GAL4 driver but no RNAi (**Figure 4A-B**). Total DAPI intensity per nucleus was quantified and normalized to control values (**Figure 4B**). *Nup98-96*-depleted salivary gland nuclei had a mean total DAPI intensity of 0.158179 relative to control salivary glands. Assuming that control nuclei have a ploidy of 1,024C [57], *Nup98-96*-depleted salivary gland nuclei had an average ploidy of ∼162C (**Figure 4B**). Consistent with our findings in S2 cells, there was a significant reduction in the percentage of *Nup98-96-*depleted salivary gland cells in S phase relative to control salivary gland cells (p<0.0001; **Supplemental Figure 4C**). Thus, we conclude that *Nup98-96-*depleted salivary glands had smaller tissue because the cells were not entering S phase, and that *Nup98-96* promotes entry into S phase in endocycling Drosophila larval salivary glands.

**Figure 4:**
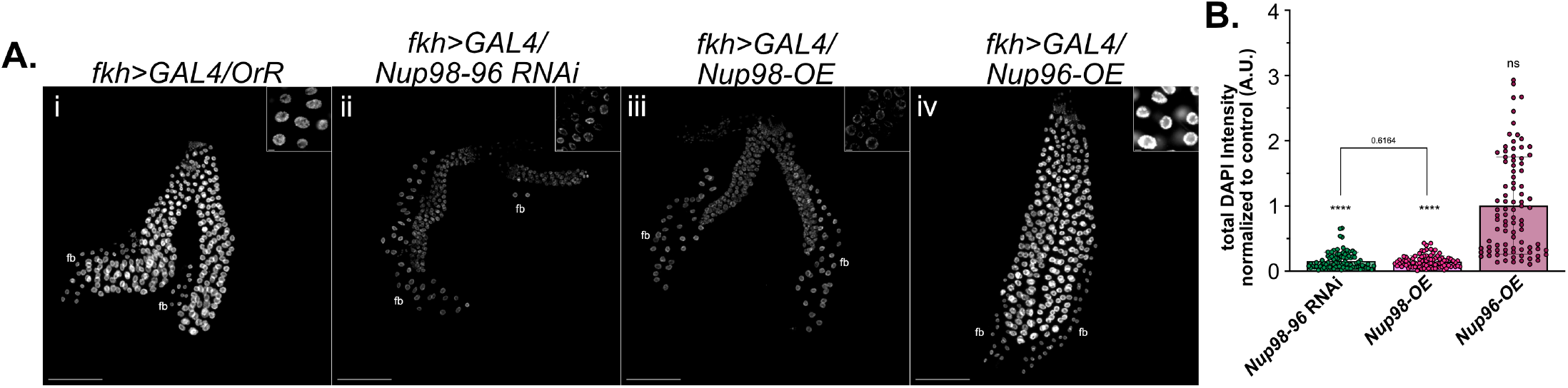
Nup98 regulates entry into S phase in Drosophila salivary glands. **(A)** Representative images of DAPI-stained 96-99 hr AEL salivary glands. 10x scale bars represent 200 µm and 60x scale bars represent 10 µm. **(Ai-iv)** 10x image of *fkh>GAL4/OrR* control, *fkh>GAL4/Nup98-96 RNAi,* f*kh>GAL4/Nup98-OE*, and *fkh>GAL4/Nup96-OE* salivary glands with inset representative 60x image. Fat bodies are labeled (fb) and genotypes are noted. **(B)** Quantification of the total DAPI intensity in arbitrary units (A.U.) per nucleus normalized to OrR control nuclei. Statistical significance was determined via One-Way ANOVA with *post-hoc* Dunnett’s test (OrR as control; n=3, N=30 nuclei per replicate; **** p<0.0001; ns=not significant). Statistical significance between *Nup98-96 RNAi* and *Nup98-OE* was determined via Student’s T-test. Error bars represent SD.

Previous studies of Nup98-96 in transcriptional regulation demonstrated that overexpression of *Nup98*, not *Nup96*, phenocopies *Nup98-96* depletion [8,9]. This dominant-negative phenotype of *Nup98* overexpression provides the opportunity to separate the function of *Nup98* and *Nup96* in S phase entry. Thus, we independently overexpressed *Nup98* and *Nup96* in larval salivary glands and measured ploidy and nuclear volume at 96-99hr (AEL). *Nup98* overexpression (*Nup98-OE*) salivary gland nuclei had a mean DAPI total intensity of 0.149652 relative to control glands that translates to an average ploidy of ∼153C (p<0.0001; **Figure 4D**). In contrast, *Nup96* overexpression (*Nup96-OE*) salivary gland nuclei had a mean DAPI total intensity of 1.00738 relative to control glands, which translates to an average ploidy of ∼1,031C (**Figure 4D**). Importantly, *Nup98-OE* salivary gland nuclei were statistically indistinguishable from *Nup98-96 RNAi* nuclei (**Figure 4D**). We observed similar results when assessing the nuclear volumes (**Supplemental Figure 4D**). Given that *Nup98* overexpression phenocopies *Nup98-96* depletion in S phase entry, we conclude that Nup98, not Nup96, promotes entry into S phase.

Previous work in Drosophila larval wing discs identified that depletion of *Nup98-96* increased entry into S phase [58]. Given that we observed decreased entry into S phase when *Nup98-96* is depleted in S2 cells and in salivary glands, we tested the same *Nup98-96 RNAi* fly line (*Nup98-96 RNAi* (BDSC)) used in the previous study to determine if the phenotypes we observe are unique to RNAi lines. Consistent with Pulianmackal et al, we observed a modest increase in DAPI total intensity and nuclear volume (**Supplemental Figure 4E-G)**. To test the efficacy of both *Nup98-96 RNAi* lines, we measured *Nup98-96* transcript levels by qPCR in salivary glands dissected from wandering third instar larvae. We found that the BDSC *Nup98-96 RNAi* line used in Pulianmackel et al., did not deplete *Nup98-96* in larval salivary glands (**Supplemental Figure 4H**). In contrast, the VDRC *Nup98-96 RNAi* line used in our work significantly reduced *Nup98-96* transcript levels (p<0.01; **Supplemental Figure 4H**). Thus, the reduced entry into S phase upon *Nup98-96* depletion in larval salivary glands correlates with reduced *Nup98-96* transcript levels.

### Nup98 regulates entry into S phase in Drosophila wing discs

Given that Nup98 regulates entry into S phase in Drosophila S2 cells and in larval salivary glands, we next asked if Nup98 regulates entry into S phase in larval wing discs. Wing discs exhibit asynchronous mitotic cellular divisions throughout larval development [59–61]. We depleted *Nup98-96* in the pouch region of 117-120hr AEL wing discs using *nub*>GAL4 and quantified the percentage of cells in S phase (**Figure 5A-B**). *nub*>GAL4 is expressed in the pouch, thus we quantified the percentage of cells in S phase in both the pouch and an unaffected control region in the notum within the same wing disc. *nub*>GAL4 *Nup98-96-*depleted wing discs had significant reductions in the ratio of cells in S phase in the pouch relative to the notum (ratio=0.034098; p<0.0001; **Figure 5B**), whereas the ratio of S phase cells in pouch relative the notum was ∼1 in control wing discs (control ratio=0.924332; **Figure 5B**). This altered ratio was due to a decrease in cells in S phase within the pouch relative to the notum (**Supplemental Figure 5A**). *Nup98-96-*depleted wing discs had irregular nuclear morphology in the pouch where DAPI staining appeared ring-like and nuclei had large chromocenters (**Figure 5Aii-ii’’**). These pouches also had a significant reduction in the number of nuclei relative to notums compared to control (control ratio=1.35156; *Nup98-96*-depleted ratio=0.703593; p<0.001; **Supplemental Figure 5B-C**). To determine if this reduction in nuclei was caused by a lack of proliferation or by apoptosis, we stained *Nup98-96-*depleted and control wing discs for the apoptotic marker cleaved-DCP1 [62]. We did not observe robust pouch-wide staining, suggesting that reduction in nuclei count is likely due to reduced proliferation rather than apoptosis (**Supplemental Figure 5D**).

**Figure 5:**
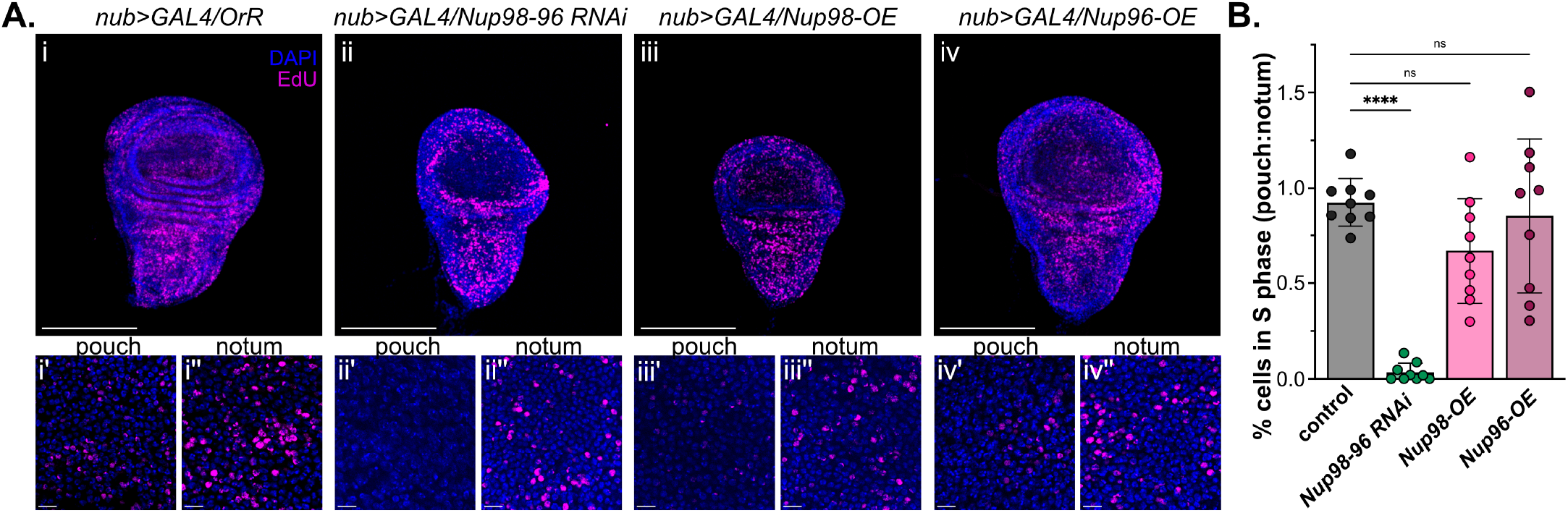
Nup98 regulates entry into S phase in Drosophila wing discs. **(A)** Representative images of DAPI-stained and EdU-pulsed 117-120 hr AEL wing discs. **(Ai-iv)** Representative images of DAPI-stained and EdU-pulsed 117-120 hr AEL wing discs. **(Ai-iv)** 10x image of *nub>GAL4/OrR* control, *fkh>GAL4/Nup98-96 RNAi*, f*kh>GAL4/Nup98-OE*, and *fkh>GAL4/Nup96-OE* wing discs. DAPI (blue) and EdU (pink) are shown. Scale bars represent 200 µm. **(Ai’-iv’)** Representative 60x images of the pouch region of each disc shown above at 10x. Scale bars represent 10 µm. **(Ai”-iv”)** Representative 60x images of the notum region of each disc shown above at 10x. Scale bars represent 10 µm. **(B)** The ratio of the percentage of cells in S phase in pouches relative to notums. Statistical significance was determined via One-Way ANOVA with *post-hoc* Dunnett’s test (*nub>GAL4/OrR* as control; n=3, N=3 discs per biological replicate; **** p<0.0001, ns=not significant). Error bars represent SD.

We next asked if *Nup98*-overexpression has a dominant-negative effect on entry into S phase in larval wing discs. To test this, we independently overexpressed *Nup98* and *Nup96* in wing disc pouches using the *nub*>GAL4 driver and measured the percentage of cells in S phase at 117-120hr AEL (**Figure 5C-D**). Overexpression of *Nup96* did not affect the ratio of S phase cells in pouch relative to the notum (ratio=0.852684; **Figure 5D**). Despite normal entry into S phase, *Nup96-OE* pouches had significantly reduced nuclei count in the pouch relative to notum (p<0.01, **Supplemental Figure 5B**). When we overexpressed *Nup98* in wing disc pouches, there was a slight reduction in the ratio of S phase cells in the pouch relative to the notum, but this reduction was not statistically significant (ratio=0.669943, **Figure 5D**). We conclude that while *Nup98-*overexpression can act as a dominant-negative block to Nup98 function, this phenotype likely depends on developmental and functional contexts.

## DISCUSSION

We previously demonstrated that *Nup98-96* is necessary for proper entry into S phase, however, the underlying mechanism was not explored [29]. In this work, we show that Nup98 promotes entry into S phase likely by regulating the expression of key cell cycle genes. First, we found that a subset of Nups, but not NPCs, affects entry into S phase, with Nup98 being the only dynamic Nup to affect the G1/S transition. Second, we show that reduced entry into S phase upon *Nup98-96* depletion corresponds to decreased transcriptional activation of cell cycle and S phase genes, including canonical E2f1 targets [34,35]. Third, we show that transcriptomic profiles of *Nup98-96*-depleted and *E2f1*-depleted cells are similar and many transcripts depend on both Nup98-96 and E2f1 for expression. Critically, we found that *Nup98-96* regulates entry into S phase regardless of cell type, tissue type, or whether cells were completing endocycles or mitotic cell cycles. Given that the overexpression of *Nup98*, which interferes with Nup98 function [8,9] phenocopies *Nup98-96* depletion, it is highly likely that Nup98, and not Nup96, is needed for S phase entry. Taken together, our observations are consistent with a model where Nup98 functions to transcriptionally promote cell cycle gene expression during the G1/S transition.

Contrary to our findings, previous work in Drosophila suggested that *Nup98-96* is required to promote or maintain a developmentally programmed G1 arrest [58]. While it is unclear why our observations are different than this previous study, our work is consistent with previous observations showing that Nup98 promotes expression of developmental and cell cycle-regulated transcripts in Drosophila Kc167 cells [5]. One reason for this discrepancy could be the RNAi lines used. We could not deplete *Nup98-96* using the same RNAi line used in this study in larval salivary glands. While there may be tissue-specific regulation or driver dependencies, we have shown that depleting *Nup98-96* with multiple independent interfering RNAs inhibits S phase entry and that Nup98-96-dependent S phase entry was seen across multiple cell and tissue types.

Drosophila Nup98 has been shown to bind to the promoter regions of active genes to regulate gene expression by several different mechanisms [4,8,9]. These include regulation of chromatin architecture through an interaction with architectural proteins [4,9], interacting with transcription factor complexes [8], affecting enhancer-promoter contacts [9], and affecting transcriptional memory [15]. While we find that *Nup98-96*-depleted S2 cells display reduced transcript levels of key cell cycle and S phase genes, and that co-depleting *dap* rescues entry into S phase, we do not know the specific molecular mechanism(s) by which Nup98 regulates transcriptional activation during the G1/S transition.

There are multiple ways Nup98 may function alongside E2F machinery to promote successful entry into S phase. It is possible that Nup98 affects recruitment of E2f1 to its transcriptional targets. Nup98 may affect local chromatin architecture to make specific genes more accessible to E2f1 for binding or could bind directly to E2f1 to promote E2f1-chromatin association. As an FG-repeat Nup with a large intrinsically disordered region (IDR), Nup98 can undergo phase separation and is known to form condensates *in vivo* and *in vitro* [18,26–28,63–66]. Further, the FG domain of Nup98 is critical for Nup98-oncofusion protein functions in acute myeloid leukemias (AML) [23–28]. Thus, it is possible that the ability of Nup98 to self-interact and phase separate affects local chromatin architecture or the ability of Nup98 to interact with E2f1 in a way that promotes E2f1 binding to chromatin.

If Nup98 helps recruit E2f1 to chromatin, either directly or indirectly, this would explain the rescue of entry into S phase that we observe when we co-deplete *Nup98-96* and *dap*, a phenotype we also observed with *E2f1* and *dap* co-depletion. CycE/CDK2 activation is the critical step for completing the G1/S transition [33,34]. Therefore, it is possible that depleting *dap* increases CycE/CDK2 activity beyond a threshold that propels cells into S phase even when *CycE* and *CDK2* expressions are dampened. A model involving insufficient CycE/CDK2 activity would also explain the lack of rescue when we co-deplete *Nup98-96* and *Rbf* or *E2f1* and *Rbf*, as there is likely not enough *CycE* transcriptional activation to complete the G1/S transition. As an alternative to promoting the E2f1 transcriptional program, Nup98 may bind directly to CycE and promote its activity. CycE/CDK2 are partly responsible for the hyperphosphorylation of Rbf in late G1 [31,67,68], thus activating E2f1, which in turn activates more *CycE* expression. Therefore, increased CycE activity would allow for both more *CycE* gene expression and successful entry into S phase. Nup98 may also compete with Dacapo for binding to CycE/CDK2, thus protecting CycE/CDK2 from degradation. This could also explain the rescue of entry into S phase that we observe when *Nup98-96* and *dap* are co-depleted. Understanding the specific mechanism(s) by which Nup98 regulates transcriptional activation of cell cycle genes will provide valuable insight into how the G1/S transition is controlled and could have implications for Nup98 oncofusions in AML pathogenesis.

## MATERIALS AND METHODS

### Drosophila cell culture

*Drosophila melanogaster* S2 cells were obtained from Drosophila Genomics Resource Center (stock number 181). Cells were maintained in log phase at 25°C in Schneider’s medium (Gibco #21720-024) supplemented with 10% heat-inactivated fetal bovine serum (GeminiBio #100-108) and 1% penicillin-streptomycin (Gibco #15140-122).

### Fly lines

Drosophila genetic stocks used in this study include Bloomington stocks 78060 (*fkh>GAL4*), 86108 (*nub>GAL4*), 64026 (*UAS-dap RNAi*), 28562 (*UAS-Nup98-96 RNAi*), 92790 (*UAS-Nup96.myc*), and 92791 (*UAS-Nup98.myc*) and Vienna Drosophila Resource Center 31198 (*UAS-Nup98-96 RNAi*). OregonR (OrR) flies were as a wild-type control.

### RNA interference

dsRNA was generated using Hi-Scribe T7 polymerase (NEB #E2050S) using primers designed with flyrnai.org/up-torr/ [69]. A list of primers used to generate dsRNA can be found in Supplemental Table S3. S2 cells were diluted to 1.5×10^6^ cells/mL in serum-free media. Cells were incubated with 20 μg of dsRNA for 45 min at 25°C. After 45 min, medium containing serum was added to cells. Cells were incubated for 3 or 5 days at 25°C. RNAi treatment durations were chosen for effective knockdown without inducing cell death. For cells with dual RNAi treatments, cells were harvested two days after initial RNAi treatment, and the RNAi process was repeated. Cells were incubated for three more days at 25°C. To confirm depletions, 1×10^6^ cells were harvested and lysed in TRIzol reagent (Invitrogen #15596026) or 2x Laemmli sample buffer (BioRad #1610737) with 50 mM DTT (Sigma Aldrich #D9779-5G). Depletions were confirmed by SDS-PAGE followed by western blotting for Elys and Nup98, or by qPCR for all other depletions.

### Western Blotting

Depletions of Elys and Nup98 were confirmed by SDS-PAGE followed by western blotting using anti-Elys [29] or anti-Nup98 [4]. Briefly, samples were boiled at 95°C and loaded onto a 4-15% Mini-PROTEAN TGX Stain-Free Gel (BioRad # 4568086). After electrophoresis, the gel was activated and imaged using a BioRad ChemiDoc^TM^ MP Imaging System. A Trans-Blot Turbo Transfer System (BioRad) was used to transfer protein to a low fluorescence PVDF membrane. Membranes were blocked with 5% non-fat milk in TBS-T (140 mM NaCl, 2.5 mM KCl, 50 mM Tris HCL pH 7.4, 0.1% Tween 20). Blots were incubated overnight at 4°C with rabbit anti-Elys (1:500) or rabbit anti-Nup98 (1:500) antibodies. Blots were washed with TBS-T and incubated with HRP-conjugated anti-rabbit secondary antibody (1:1000; Jackson ImmunoResearch # 111-035-003). Blots were washed again, incubated in Clarity^TM^ Western ECL Substrate (BioRad #180-5061) following manufacturer’s recommendations, and imaged on a BioRad ChemiDoc^TM^ MP Imaging System.

### Quantitative PCR

Total RNA was isolated from 1.0×10^6^ S2 cells or wandering third instar larval salivary glands using 1 mL of TRIzol reagent (Invitrogen #15596026). RNA was DNase-treated (NEB #M0303S), purified by phenol-chloroform extraction, and 50 ng of DNA-free RNA was used for one-step cDNA synthesis with iScript gDNA Clear cDNA synthesis kit (BioRad #1725035). Quantitative real-time PCRs (qPCRs) were carried out on the resulting cDNA to measure mRNA levels for target genes using a BioRad CFX Maestro. qPCR primers were designed using flyrnai.org/flyprimerbank [69] (for primer list, see Supplemental Table S4). All mRNA levels were normalized to *GAPDH1* transcript levels. Relative transcript levels (ΔΔCq) and fold-changes (2^-ΔΔCq^) were calculated compared to non-targeting *GFP* RNAi control samples. Statistical significance was determined via One-Way ANOVA with *post-hoc* Dunnett’s test with non-targeting *GFP* RNAi as control on GraphPad PRISM (version 10.1.1; GraphPad Software).

### Flow cytometry

Cell cycle profiles were generated as previously described [29]. Briefly, 10×10^6^ RNAi-treated cells were pulsed with 20 µM EdU (ChemCruz #sc-284628A) for 20 min. Harvested cells were washed with 1xPBS and fixed overnight with ice-cold 70% ethanol. Fixed cells were washed with 1xPBS and then permeabilized with 1xPBX (1xPBS with 0.1% Triton X-100) for 1 hr at room temperature. Incorporated EdU was click-labeled with AlexaFluor 555 Azide (Invitrogen #A20012) by incubating cells with 4 mM CuSO_4_, 6.25 µM AlexaFluor 555 Azide, and 2 mg/mL sodium ascorbate in 1xPBS for 30 min at room temperature. Cells were washed with 1xPBX and stained overnight at 4°C with 10 µg/mL DAPI (Roche # 1026276001) in 1xPBX. DNA content and EdU intensity were assessed using a 5-laser Fortessa analytical flow cytometer. Wild-type cells were used for compensation controls. At least three biological replicates were used per sample. Flow cytometry data was plotted and analyzed using FlowJo (version 10.10.0; BD Biosciences). For an example of gating used in these experiments, see Figure S1A. Statistical significance was determined via One-Way ANOVA with *post-hoc* Dunnett’s test with non-targeting *GFP* RNAi as control on GraphPad PRISM.

### Immunofluorescence

To measure the percentage of cells in S phase, RNAi cells were pulsed with 20 µM EdU for 20 min. 1×10^6^ cells per sample were harvested and attached to Concanavalin A (FisherScientific # AAJ61221MC)-coated coverslips for 15 min at 25°C. Cells were washed with 1xPBS and fixed with 4% PFA for 15 min at room temperature. Fixed cells were click-labeled as previously described (see flow cytometry) with AlexaFluor 488 Azide (Invitrogen #A10266). Cells were washed with 1xPBX and stained for 20 min at room temperature with 10 µg/mL DAPI in 1xPBX. Coverslips were washed once with 1xPBX and mounted with VectaShield mounting media (Vector Labs #H-1900-10). Slides were imaged at 60x on a Nikon Eclipse Ti microscope. Three biological replicates per sample. To quantify the EdU signal, Nikon NIS Elements software was used to generate regions of interest (ROIs) using DAPI signal. Mean FITC (AlexaFluor 488) intensity for each ROI was determined for at least 500 nuclei per biological replicate. Nuclei were binned into EdU-negative and EdU-positive groups based on mean FITC signal, and thresholding between EdU-negative and EdU-positive signals was defined using wild-type cells that were not pulsed with EdU but otherwise underwent the same processing as EdU-pulsed cells. The percent of cells in S phase was quantified for each biological replicate, and statistical significance was determined via One-Way ANOVA with *post-hoc* Dunnett’s test with non-targeting *GFP* RNAi as control on GraphPad PRISM.

To assess nuclear pore complex integrity, cells were pulsed with EdU, plated on Concanavalin A-coated coverslips, fixed, and permeabilized as described above. After permeabilization, cells were blocked in 1xPBX supplemented with 1% bovine serum albumin (BSA) (Fisher Bioreagents #BP9706) and 0.2% Normal Goat Serum (Sigma Aldrich #G9023) for 1 hr at room temperature. Cells were click-labeled as previously described (see flow cytometry) with AlexaFluor 488 Azide (Invitrogen #A10266). Cells were washed with 1xPBX. After washing, cells were incubated in anti-mAb414 (1:300; Abcam #ab24609) for 2 hr at room temperature. Cells were washed again with 1xPBX. Cells were incubated in 568-conjugated anti-mouse secondary antibody (1:500; Invitrogen #A-11031) for 2 hr at room temperature. Cells were washed with 1xPBX and stained for 20 min at room temperature with 10 µg/mL DAPI in 1xPBX. Cells were washed with 1xPBX and mounted with VectaShield mounting media (Vector Labs #H-1900-10). Slides were imaged at 60x on a Nikon Spinning Disk confocal microscope for three biological replicates per sample. The same laser intensities and exposure times for channels were used for all samples per biological replicate. To quantify mAb414 signal, 4-point ellipses were drawn to define ROIs around 50 nuclei per biological replicate using NIS-Elements AR 4.60 software. Mean Texas Red (mAb414) intensity for each ROI was quantified. Wild-type cells that were not incubated in anti-mAb414 primary antibody but otherwise underwent the same processing as mAb414-stained cells were used as negative control. Mean mAb414 intensity per nucleus was plotted and statistical significance was determined via Student’s t-test on GraphPad PRISM.

### Salivary gland immunofluorescence

To assess relative ploidy and fraction of cells in S phase, male flies from RNAi lines were crossed to *fkh*>GAL4 virgin females. Flies were fed dry yeast and independent crosses were set up for three biological replicates per genotype. Salivary glands were dissected from 95-98 AEL larvae in 1xPBS and pulsed with 200 µM EdU for 10 min at room temperature. Glands were fixed in 4% PFA for 30 min at room temperature. After fixation, glands were washed three times with 3xPBX (1xPBS supplemented with 0.3% TritonX-100) for 15 min each to allow for PFA removal and tissue permeabilization. EdU was click-labeled with Alexafluor 488 Azide or AlexaFluor 647 (Invitrogen #10277) as described above. To optimally DAPI stain glands to quantify ploidy, glands were incubated in 50 ng/mL DAPI at 4°C for 1 hr and washed overnight at 4°C with 3xPBX [56,70]. Glands were mounted in VectaShield mounting media. Glands were imaged at 10x and 60x on a Nikon Eclipse Ti microscope. The same laser intensities and exposure times for channels were used for all samples per biological replicate. For 10x images, z-slices were acquired every 5 µm for the total height of the glands. For 60x images, z-slices were acquired every 1 µm for the total height of the glands. DNA content was analyzed by quantifying total DAPI intensity per nucleus and nuclear volume using ilastik software [71]. Data was quantified for 10 nuclei per gland from 3 separate glands per biological replicate. Total DAPI intensity per nucleus was normalized to OrR control nuclear values. Statistical significance was determined via One-Way ANOVA with *post-hoc* Dunnett’s test with OrR as control on GraphPad PRISM. When comparing *Nup98-96 RNAi* normalized DAPI intensity and *Nup98-OE* normalized DAPI intensity, statistical significance was determined via Student’s t-test on GraphPad PRISM. When comparing *Nup98-96 RNAi* (BDSC) DAPI intensity and volumes to OrR control, statistical significance was determined via Student’s t-test on GraphPad PRISM. The percentage of cells in S phase was calculated by manually counting the number of OrR and *Nup98-96 RNAi* nuclei and EdU-positive nuclei per lobe using NIS-Elements AR (version 4.60.00 software; Nikon Instruments, Inc.). Statistical significance was determined via Student’s t-test on GraphPad PRISM. Scale bars were added to images using FIJI (version x86-64; [72]).

### Wing disc immunofluorescence

To assess the fraction of cells in S phase in larval wing discs, male flies from RNAi lines were crossed to *nub*>GAL4 virgin females. Flies were fed dry yeast and independent crosses were set up for three biological replicates per genotype. Wing discs were dissected from 117-120hr AEL larvae in 1xPBS and pulsed with 50 µM EdU for 10 min at room temperature. Wing discs were fixed in 4% PFA for 30 min at room temperature. After fixation, discs were washed three times with 3xPBX (1xPBS supplemented with 0.3% TritonX-100) for 15 min each to allow for PFA removal and tissue permeabilization. EdU was click-labeled with AlexaFluor 647 Azide as described above. Discs were incubated in 50 ng/mL DAPI at room temperature for 20 min and washed with 3xPBX for 15 min. Discs were mounted in VectaShield mounting media and imaged at 10x and 60x on a Nikon Eclipse Ti spinning disc confocal microscope. 1 z-slice was acquired for 10x images. For 60x images, z-slices were acquired every 1 µm for the total height of the discs. To quantify the percentage of cells in S phase, a 4143.75 µm^2^ area of the pouch and a 4143.75 µm^2^ area of the notum of each wing disc was defined using FIJI (version x86-64; [72]). Total nuclei and the number of EdU-positive nuclei were manually counted from a single z-slice from each pouch and notum using NIS-Elements AR (version 4.60.00 software; Nikon Instruments, Inc.). The ratio of S phase cells was calculated by dividing the percentage of cells in S phase in the pouches of each wing disc to the percentage of cells in S phase from the notums. The ratio of nuclei count was calculated by dividing the number of pouch nuclei per wing by the number of notum nuclei per wing. Statistically significant differences in the ratio of S phase cells and nuclei count were determined via One-Way ANOVA with *post-hoc* Dunnett’s test with OrR as control on GraphPad PRISM.

To assess apoptosis activation, wing discs were dissected from 117-120hr AEL OrR and *Nup98-96 RNAi* larvae in 1xPBS. Discs were fixed and permeabilized as described above. Discs were incubated in blocking solution (3xPBX supplemented with 1% BSA and 0.2% Normal Goat Serum for 1 hr at room temperature. Discs were incubated in anti-cleaved DCP1 (1:500; Cell Signaling, #9578) for 2 hr at room temperature. Discs were washed with 3xPBX, then incubated in 568-conjugated anti-rabbit secondary antibody (1:500; Invitrogen #A-11011) for 1 hr at room temperature. Discs were washed with 3xPBX. Discs were incubated in 50 ng/mL DAPI at room temperature for 20 min and washed with 3xPBX for 15 min. Discs were mounted in VectaShield mounting media and imaged at 10x and 60x on a Nikon Eclipse Ti spinning disc confocal microscope. 1 z-slice was acquired for 10x images. For 60x images, z-slices were acquired every 1 µm for the total height of the discs.

### RNA-sequencing

Total RNA was isolated from 1×10^6^ N*up98-96* RNAi cells and non-targeting *GFP* RNAi cells using 1 mL TRIzol reagent. 5 µg of RNA was DNase-treated and purified by phenol-chloroform extraction as described above (see Quantitative PCR). Ribosomal RNA (rRNA) was depleted using the RiboMinus Eukaryotic Kit for RNA-Seq (Invitrogen #A1083708) following manufacturer’s recommendations. 5 ng of rRNA-free RNA per sample was used to synthesize cDNA library with the NEBNext Ultra^TM^ II RNA Library Prep Kit for Illumina (NEB #E770S) and NEBNext Multiplex Oligos for Illumina (Dual Index Primers Set 1) (NEB #E7600S) following manufacturer’s recommendations. Libraries were sequenced on an Illumina NovaSeq 6000 by Vanderbilt Technologies for Advanced Genomics (VANTAGE). For *E2f1-*depleted cells, total RNA was isolated from 1×10^6^ *E2f1* RNAi cells and non-targeting *GFP* RNAi cells using 1 mL TRIzol reagent. 5 µg of RNA was DNase-treated and purified by phenol-chloroform extraction as described above. DNA-free RNA was sent to Plasmidsaurus for poly(A) mRNA enrichment, cDNA synthesis, Illumina library preparation, and RNA-sequencing with custom analysis and annotation.

### RNA-sequencing analysis

Reads from paired-end Illumina sequencing were trimmed using –g, selected for reads at least 100 bp long with –l 100, and quality was assessed using fastp (version 0.20.1; [73]. Reads were aligned to the BDFP6.46.112 *D. melanogaster* reference genome using STAR (version 2.5.4b; [74]). The BDFP6.46.112 *D. melanogaster* reference genome and its annotations were downloaded from Ensembl (release 112). SAMtools (version 1.18; [75]) was used to convert aligned read files to bam files, and index and sort bam files. BEDTools (version 2.28.0; [76]) was used to exclude reads corresponding to *Nup98-96* dsRNA from Chr3R 23,779,890-23,780,210. Feature counts were generated using Subread (version 2.0.6; [77,78]). Differential expression analysis was done with DESeq2 (version 1.46.0; [79]). Volcano plots were made with ggplot2 (version 4.0.2; [80]) and heatmaps were made with pheatmap (version 1.0.13; [81]). Gene ontology analysis was conducted on STRING (version 12.0; Szklarczyk et al., 2022). Reads from single-end Illumina sequencing were processed as described above, with the exceptions of selecting reads at least 50 bp long with –l 50 and reads from Chr3R 23,779,890-23,780,210 were not removed. Generation of feature counts, differential expression analysis, and gene ontology analysis were conducted as described above. To identify overlap between transcripts with significantly decreased levels in both RNA-seq datasets, Venn diagrams were made with venneuler (version 1.1-4; [82]) and overlapping transcripts were identified with ggVennDiagram (version 1.5.7; [83]). Statistically significant overlap was determined via hypergeometric test [84].

Previously published data were retrieved for *lacZ* control and *DREF* RNAi S2R+ cell RNA-seq (GEO: GSE117217; [51]). Single-end reads were processed as described above, with the exceptions of selecting reads at least 50 bp long with –l 50 and reads from Chr3R 23,779,890-23,780,210 were not removed. Generation of feature counts, differential expression analysis, and gene ontology analysis were conducted as described above.

### ChIP-sequencing analysis

Previously published S2 cell ChIP-seq Nup98 binding peaks were used [9]. Peak coordinates were converted from dm3 to dm6 using LiftOver Genome annotation tool from UCSC Genome Browser [85]. Peak annotation and motif analysis were done using HOMER (version 5.1; [86]). Previously published ChIP-sequencing data from Drosophila Kc167 cells was retrieved for Nup98 (GEO: GSE80700; [45]) and corresponding IgG control (GEO: GSE63518; [87]). Reads were trimmed using –g and quality was assessed using fastp (version 0.20.1; [73]). Reads were aligned to the BDFP6.46.112 *D. melanogaster* reference genome allowing for 1 mismatch using bowtie2 (version 2.5.4; [88]). SAMtools (version 1.18; [75]) was used to convert aligned read files to bam files, index bam files, sort bam files, and merge IgG read files. Duplicate reads were marked and removed with the Picard Toolkit (version 3.1.0; Broad Institute). Coverage files were generated using Deeptools: BamCoverage (version 3.5.6; [89]). Statistically significant Nup98 peaks over IgG control were called using macs2 (version 2.2.9.1; [90,91]). Peak annotation and motif analysis were done using HOMER (version 5.1: [86]). Overlap between genes with Nup98 binding peaks in their promoters [9] and transcripts with significant differential levels after *Nup98-96* depletion in S2 cells was identified using ggVennDiagram (version 1.5.7; [83]) and visualized using venneuler (version 1.1-4; [82]). Statistically significant overlap was determined via hypergeometric test [84]. Overlap between Nup98 binding peaks in their promoters [9], transcripts with significantly downregulated levels after *Nup98-96* depletion in S2 cells, and transcripts with significantly downregulated levels after *E2f1* depletion in S2 cells was identified using ggVennDiagram (version 1.5.7; [83]) and visualized using UpSetR (version 1.4.0; [92]). Statistically significant overlap was determined via multivariate hypergeometric test [84].

## ACKNOWLEDGMENTS

Flow Cytometry experiments were performed in the Vanderbilt Flow Cytometry Shared Resource that is supported by the Vanderbilt Ingram Cancer Center (P30 CA068485) and the Vanderbilt Digestive Disease Research Center (DK068404). Illumina sequencing was performed at the Vanderbilt Technologies for Advanced Genomics (VANTAGE) core. Confocal microscopy experiments were performed in part through the Vanderbilt Cell Imaging Shared Resource that is supported by NIH grants, CA68485, DK58485, and EY08126. We thank Anneliese Schroer for providing feedback on the manuscript. We thank Jianhao Li for assistance in dissecting wing discs. This research was supported by NIH grants 2R35GM128650 to J.T.N and 2R01GM124143 to M.C.

## AUTHOR CONTRIBUTIONS

Conceptualization, E.M.M., J.T.N, and M.C.; formal analysis, E.M.M. and A.S.; investigation, E.M.M.; resources, M.C.; writing – original draft, E.M.M and J.T.N.; writing – review & editing, E.M.M., J.T.N., and M.C.; visualization, E.M.M.; supervision, J.T.N.; project administration, J.T.N.; funding acquisition, J.T.N.

