## Supplemental material for "Nup98 regulates the G1/S transition"

### Supplemental Figure 1

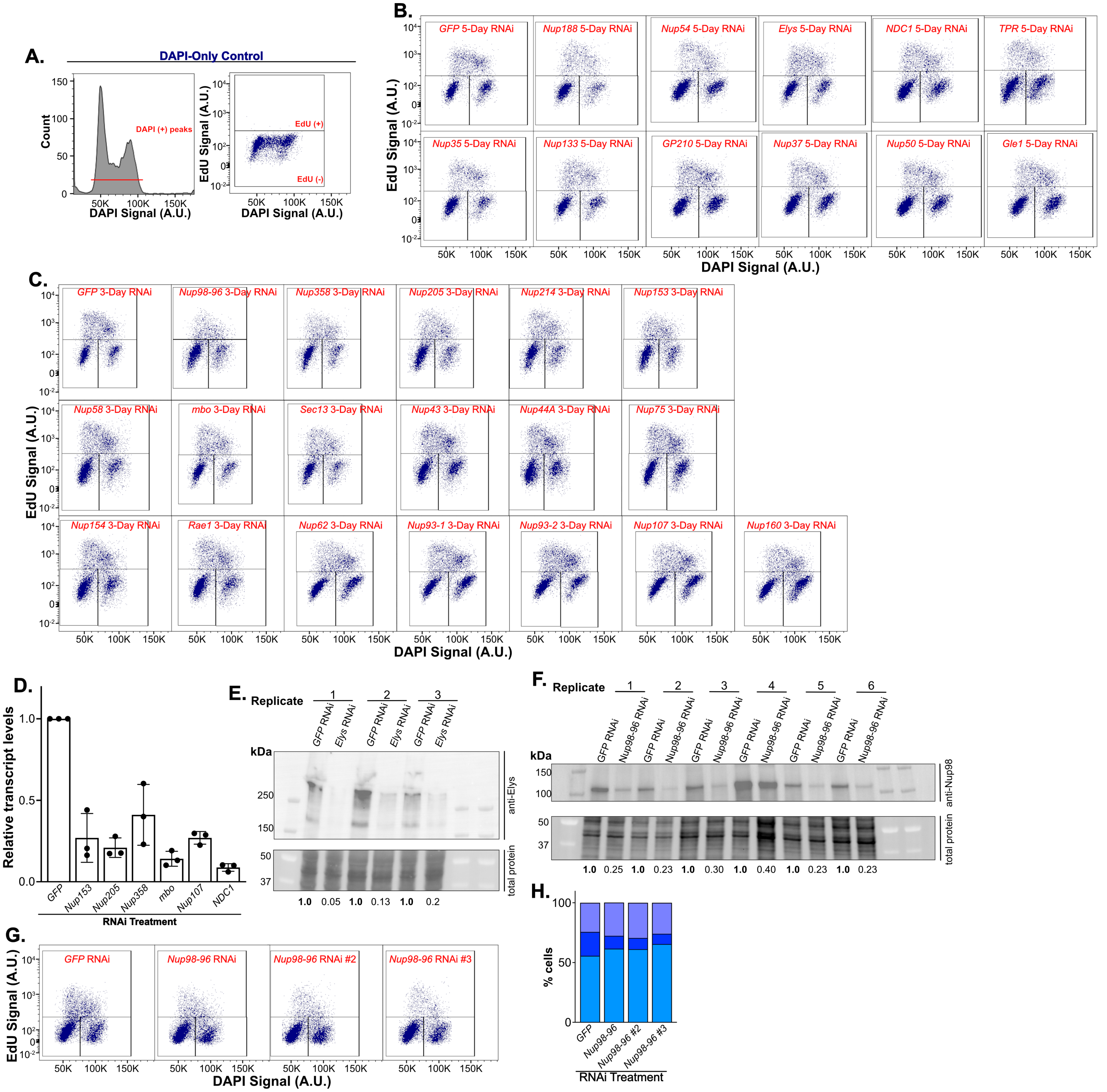

**Supplemental Figure 1: A subset of Nups impacts the G1/S transition. (A)** Representative plots on DAPI-stained S2 cells showing gating strategy. (left) gating of DAPI-positive signal. (right) gating of EdU-positive and EdU-negative signal was done using cells that were DAPI-stained but not pulsed with EdU. **(B-C)** Representative horseshoe plots were generated by plotting DNA content (DAPI) against EdU signal. **(B)** Representative plots from 5-day control RNAi (*GFP* RNAi) and 5-day *Nup* RNAi cells are shown. Gates indicating G1, S, and G2/M populations are shown (black boxes) and RNAi targets are noted in red. **(C)** Representative plots from 3-day control RNAi (*GFP* RNAi) and 3-day *Nup* RNAi cells are shown. Gates indicating G1, S, and G2/M populations are shown (black boxes) and RNAi targets are noted in red. **(D)** transcript levels relative to *GAPDH1* levels for control *GFP* RNAi, *Nup153* RNAi, *Nup205* RNAi, *Nup358* RNAi, *mbo* RNAi, *Nup107* RNAi, and *NDC1* RNAi respective knockdowns were measured by qPCR and normalized to control (n=3). Error bars represent SD. **(E)** *Elys* RNAi was verified by western blotting using anti-Elys. Values below each lane represent Elys band volume normalized to total lane protein. **(F)** *Nup98-96* RNAi was verified by western blotting using anti-Nup98. Values below each lane represent Nup98 band volume normalized to total lane protein. **(G)** Horseshoe plots were generated by plotting DNA content (DAPI) against EdU signal for non-targeting *GFP* control RNAi and *Nup98-96* RNAi driven by 3 independent dsRNAs. RNAi targets are noted in red. **(H)** The percentages of cells in G1 (light blue), S (blue), and G2/M (purple) phases were quantified.

#### Supplemental Figure 2

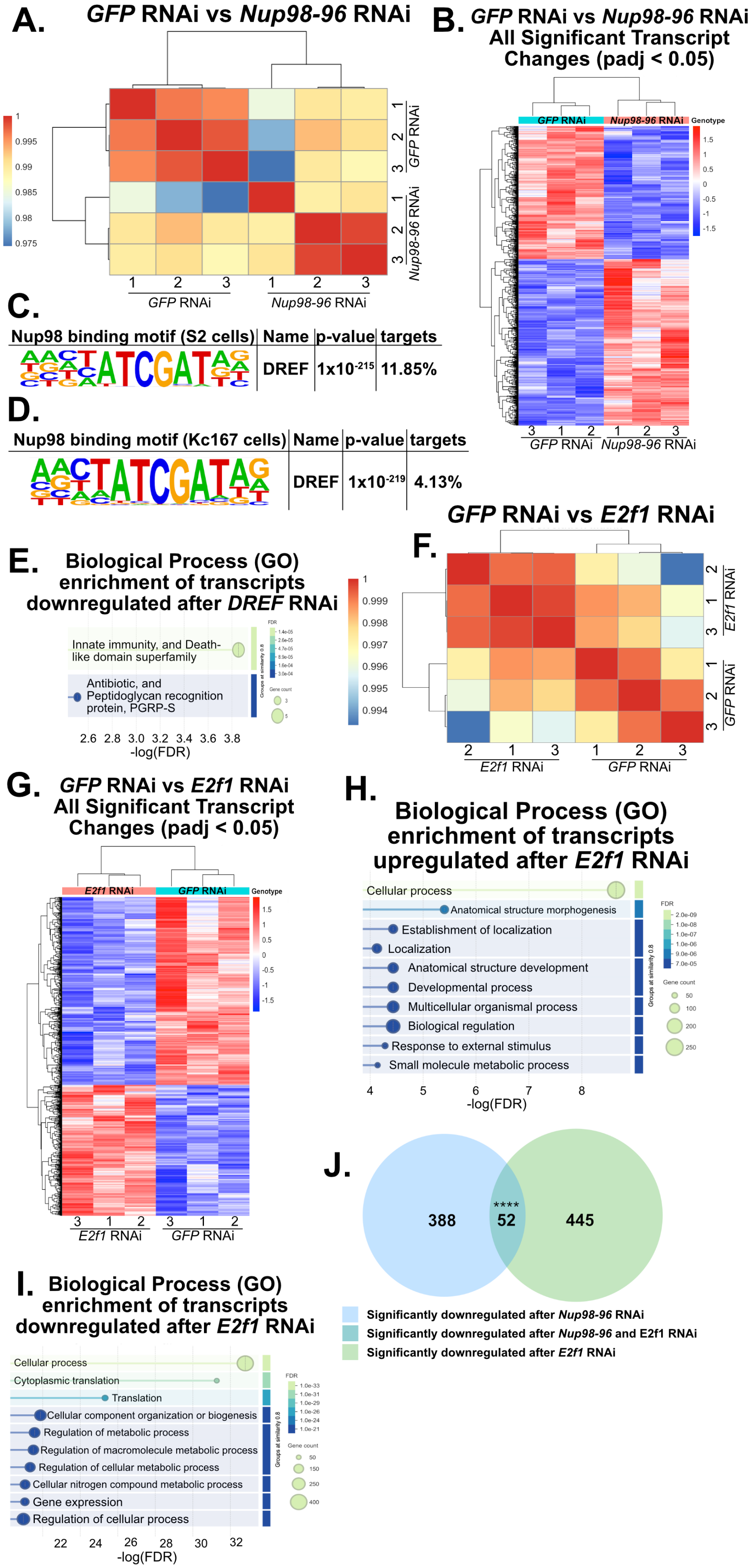

**Supplemental Figure 2: *Nup98-96* regulates the transcriptional activation of cell cycle-related genes and E2f target genes. (A)** Correlation matrix representing clustering of control *GFP* RNAi and *Nup98-96* RNAi samples. Color scale indicates Pearson coefficient. Numbers represent biological replicate. **(B)** Heatmap representing clustering of *GFP* and *Nup98-96* RNAi samples based on significant differentially expressed transcripts. Color scale indicates z-score. Numbers represent biological replicate **(C)** HOMER visualization of Nup98 binding motif corresponding to DREF motif enrichment in S2 cells. **(D)** HOMER visualization of Nup98 binding motif corresponding to DREF motif enrichment in Kc167 cells. **(E)** Gene ontology analysis of significantly downregulated transcripts after *DREF*-depletion sorted by FDR. Color scale indicates FDR and circle size represents transcript count. **(F)** Correlation matrix representing clustering of control *GFP* RNAi and *E2f1* RNAi samples. Color scale indicates Pearson coefficient. Numbers represent biological replicate. **(G)** Heatmap representing clustering of *GFP* and *E2f1* RNAi samples based on significant differentially expressed transcripts. Color scale indicates z-score. Numbers represent biological replicate. **(H)** Gene ontology analysis of significantly upregulated transcripts after *E2f1*-depletion sorted by FDR. Color scale indicates FDR and circle size represents transcript count. **(I)** Gene ontology analysis of significantly downregulated transcripts after E2f1-depletion sorted by FDR. Color scale indicates FDR and circle size represents gene count. **(J)** Venn diagram representing overlap between genes significantly downregulated after *Nup98-96*-depletion and *E2f1*-depletion. Numbers represent transcript count. Statistical significance was determined via hypergeometric test. **** $p<0.0001$.

#### Supplemental Figure 3

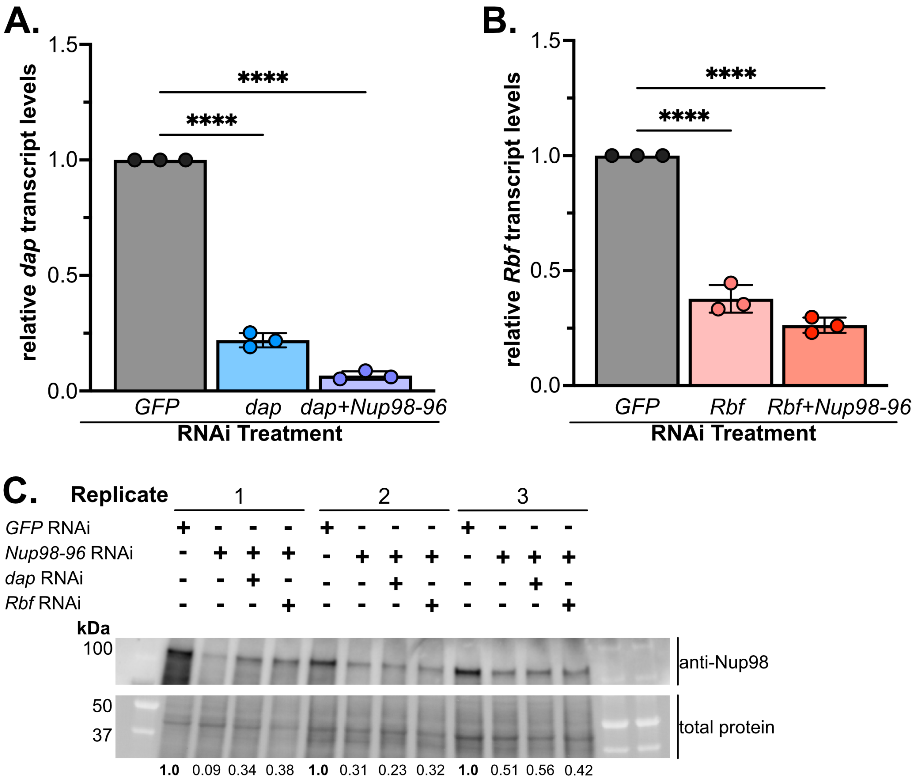


**Supplemental Figure 3: *Nup98-96* and *dacapo* have opposing functions in regulating the G1/S transition. (A)** *dap* transcript levels relative to *GAPDH1* levels were measured by qPCR for control *GFP* RNAi, *dap* RNAi, and *dap+Nup98-96* RNAi samples and normalized to control. Error bars represent SD. Statistical significance was determined via One-Way ANOVA with *post-hoc* Dunnett's test (*GFP* RNAi as control; n=3; **** p<0.0001). **(B)** *Rbf* transcript levels relative to *GAPDH1* levels were measured by qPCR for control *GFP* RNAi, *Rbf* RNAi, and *Rbf+Nup98-96* RNAi samples and normalized to control. Error bars represent SD. Statistical significance was determined via One-Way ANOVA with *post-hoc* Dunnett's test (*GFP* RNAi as control; n=3; **** p<0.0001). **(C)** *Nup98-96* RNAi was verified by western blotting using anti-Nup98. Values below each lane represent Nup98 band volume normalized to total lane protein.

#### Supplemental Figure 4

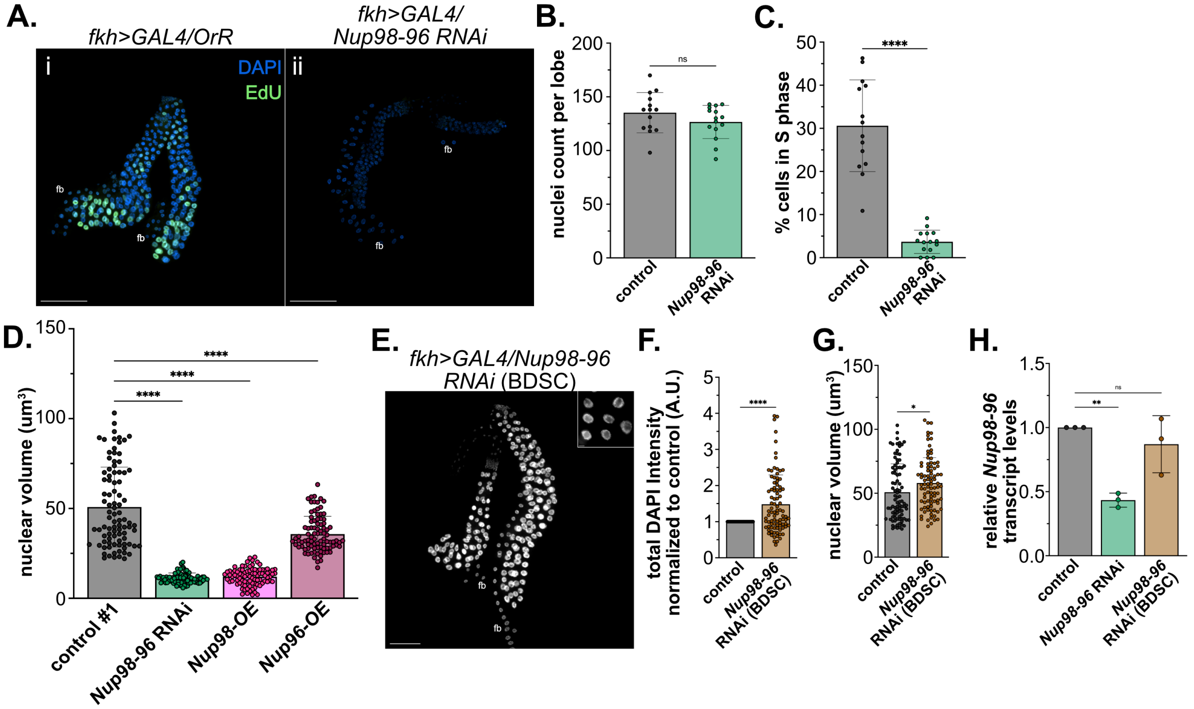


**Supplemental Figure 4: Nup98 regulates entry into S phase in Drosophila salivary glands.** **(A)** Representative 10x images of *fkh>GAL4/OrR* control and *fkh>GAL4/Nup98-96 RNAi* salivary glands that were pulsed with EdU (green) and stained with DAPI (blue). Scale bars represent 200 µm. **(B)** The number of nuclei per lobe was manually counted for OrR control (n=3, N=14 lobes) and *Nup98-96 RNAi* salivary glands (n=3, N=15 lobes). Statistical significance was determined via Student's T-test (ns=not significant). Error bars represent SD. **(C)** The % of cells in S phase was calculated by manually counting the number of nuclei per lobe and the number of EdU-positive nuclei per lobe for OrR control (n=3, N=14 lobes) and *Nup98-96 RNAi* salivary glands (n=3, N=15 lobes). Statistical significance was determined via Student's T-test (**** p<0.0001). Error bars represent SD. **(D)** Nuclear volumes were calculated for two independent OrR controls, *Nup98-96 RNAi*, *Nup98-OE*, and *Nup96-OE* salivary glands. Statistical significance was determined via One-Way ANOVA with *post-hoc* Dunnett's test (OrR as control; n=3, N=30 nuclei per replicate; **** p<0.0001, ns=not significant). Error bars represent SD. **(E)** Representative 10 image of DAPI-stained 96-99 hr old *Nup98-96* (DGRC) salivary glands with inset 60x image. 10x scale bars represent 200 µm and 60x scale bars represent 10 µm. Fat bodies are labeled (fb) and genotype is noted. **(F)** Quantification of the total DAPI intensity in arbitrary units (A.U.) per nucleus normalized to OrR control nuclei. Statistical significance was determined via Student's T-test (n=3, N=30 nuclei per replicate; **** p<0.0001). Error bars represent SD. **(G)** Nuclear volumes were calculated for OrR control and *Nup98-96 RNAi* (DGRC) salivary glands. Statistical significance was determined via Student's T-test (n=3, N=30 nuclei per replicate; * p<0.05). Error bars represent SD. **(H)** *Nup98-96* transcript levels relative to *GAPDH1* were measured by qPCR and normalized to control. Statistical significance was determined via One-Way ANOVA with *post-hoc* Dunnett's test (OrR as control; n=3, ** p<0.01; ns=not significant). Error bars represent SD.

**Supplemental Figure 5**

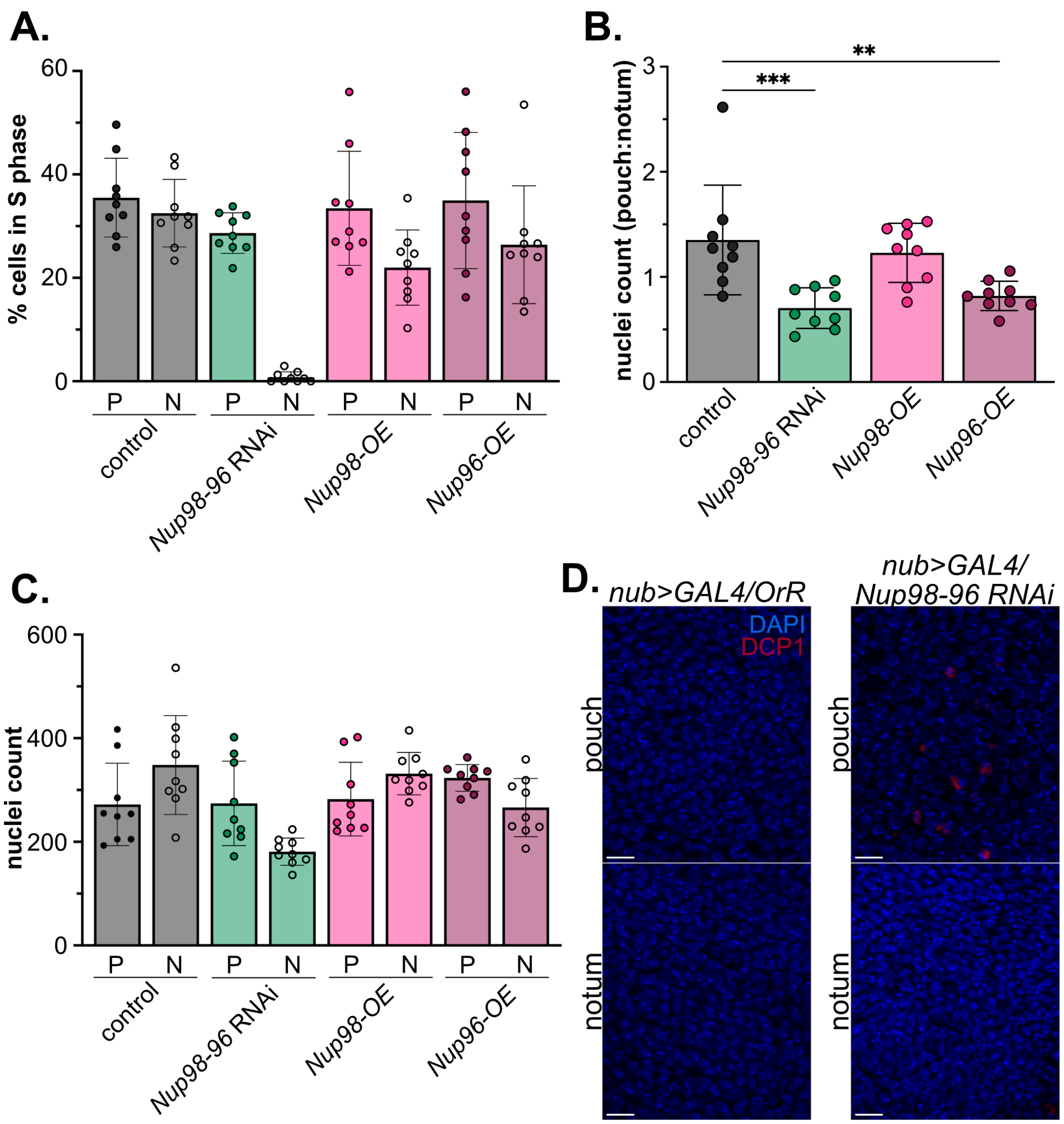


**Supplemental Figure 5: Nup98 regulates entry into S phase in Drosophila wing discs. (A)** The percentage of cells in S phase in pouches (P) and notums (N) were quantified for *nub>GAL4/OrR* control, *fkh>GAL4/Nup98-96 RNAi*, f*kh>GAL4/dap RNAi*, *fkh>GAL4/dap+Nup98-96 RNAi*, *fkh>GAL4/Nup98-OE*, and *fkh>GAL4/Nup96-OE* wing discs. n=3, N=3 discs per biological replicate. **(B)** The ratio of nuclei count in pouches relative to notums. Statistical significance was determined via One-Way ANOVA with *post-hoc* Dunnett's test (*nub>GAL4/OrR* as control; n=3, N=3 discs per biological replicate; *** $p<0.001$, ** $p<0.01$). **(C)** Nuclei count in pouches (P) and notums (N) were quantified for nub>GAL4/OrR control, *fkh>GAL4/Nup98-96 RNAi*, f*kh>GAL4/dap RNAi*, *fkh>GAL4/dap+Nup98-96 RNAi*, *fkh>GAL4/Nup98-OE*, and *fkh>GAL4/Nup96-OE* wing discs. n=3, N=3 discs per biological replicate. **(D)** Representative 60x images of *fkh>GAL4/OrR* control and *fkh>GAL4/Nup98-96 RNAi* wing disc pouches (top) and notums (bottom) that were stained for cleaved-DCP1 (red) and DAPI (blue). Scale bars represent 10 µm. n=1, N=3 wing discs.
